# Connectivity for Rapid Synchronization in a Neural Pacemaker Network

**DOI:** 10.1101/2020.10.05.325613

**Authors:** Ezekiel Williams, Aaron R. Shifman, John E. Lewis

## Abstract

Synchronization is a fundamental property of biological neural networks, playing a mechanistic role in both healthy and disease brain states. The medullary pacemaker nucleus of the weakly electric fish is a synchronized network of high-frequency neurons, weakly coupled via gap junctions. Synchrony in the pacemaker is behaviourally modulated on millisecond timescales, but how gap junctional connectivity enables such rapid resynchronization speeds is poorly understood. Here, we use a computational model of the pacemaker, along with graph theory and predictive analyses, to investigate how network properties, such as randomness and the directionality of coupling (bidirectional/non-rectifying versus directional/rectifying gap junctions) characterize the fast synchronization of the pacemaker network. Our results provide predictions about connectivity in the pacemaker and insight into the relationship between structural network properties and synchronization dynamics in neural systems more generally.

## Introduction

Synchronization between neurons is a dynamical process that underlies normative cognitive phenomena and disease states alike. In the healthy brain, synchronization of neural oscillations is believed to drive long-range communication between regions [35], enabling memory formation [27] and attentional modulation [39]. In the diseased brain, synchronization can hijack large networks resulting in epileptic seizures [24]. It follows that synchronization must be tightly regulated, balancing the coordination of neural activity with the loss of information seen when many neurons produce the same output. An important aspect of this regulation is temporal [7, 15, 28, 40]; neural systems must be able to switch between synchronized and desynchronized states in a time-efficient fashion. The mechanisms that allow such transient shifts in synchrony are not clear.

The Pacemaker Network (PN) of weakly electric fish controls the timing of the Electric Organ Discharge (EOD) produced by the fish to sense their environment [19]. Though the EOD is one of the most temporally precise oscillations known to biology [22], its dynamics are actively modulated by the fish on millisecond timescales to produce communication signals [44, 36]. Because tight spike-time synchronization and rapid-resynchronization in PN cells underlies the unique dynamic phenomena of these EOD modulations, the PN represents an intriguing model system for the study of the temporal modulation of neural synchrony.

The PN is a medullary nucleus composed of roughly 100-150 neurons [20, 21]. The majority of these cells are the gap-junctionally coupled “pacemaker cells”, intrinsic to the PN, that form a sub-network of synchronized endogenous oscillators responsible for EOD timing [21, 32, 17]. Pacemaker cells are apparently connected in a random fashion to approximately 3 to 7 neighbours [6, 21] via rectifying electrical gap junctions [20]. Interestingly, the molecular properties of rectifying gap junctions have been recently identified [30] and, while they are found in a number of systems, their functional roles are not well-understood [16]. We wish to identify the impact of such network features on PN synchronization dynamics. Indeed, graph theoretic metrics related to the nature of network connectivity can be associated with the speed and robustness of oscillator synchronization [3, 2, 10, 26, 25, 42]. As a first step towards generating testable hypotheses about PN structure-function and synchronization dynamics, we investigate how network connectivity influences speed of spike-time synchronization in a biologically-detailed model of the PN [29, 17]. Our results are divided into two main sections. First, we ask how different network parameters interact to produce the rapid synchronization times observed in the PN. Second, we ask how predictive network structure alone, rather than individual neuron properties, is in determining synchronization speed. To do this, we quantify synchronization time using the Kuromoto coefficient [1] and connectivity using graph theoretic properties and the Laplacian, a matrix related to the adjacency matrix of the network [18]. Our results suggest that the random nature of electrical connectivity in the PN leads to fast synchronization and may be an evolutionary adaptation to electrocommunication signal generation.

## Methods

We begin this section by introducing, in a biological context, the parameters from graph theory that were used to characterize PN model network structure. We then discuss the model itself, followed by data analysis methods, before proceeding to the results.

### Network Parameters

We focus on three graph-theoretic parameters in this study: (i) randomness of connectivity; (ii) directionality; (iii) degree homogeneity. These parameters were chosen in light of the suggested network properties of the PN [6, 21].

Randomness of connectivity, or network randomness, can be thought of as the extent to which a network differs from a perfectly ordered lattice on one extreme, to a more randomly connected synaptic pattern on the other [37]. We quantify network randomness by *q*, the mean ratio of randomly inserted to ordered connections in a network (Figure 1.A). It has been shown to strongly effect synchronization dynamics in a wide variety of oscillating systems [11].

**Figure 1:**
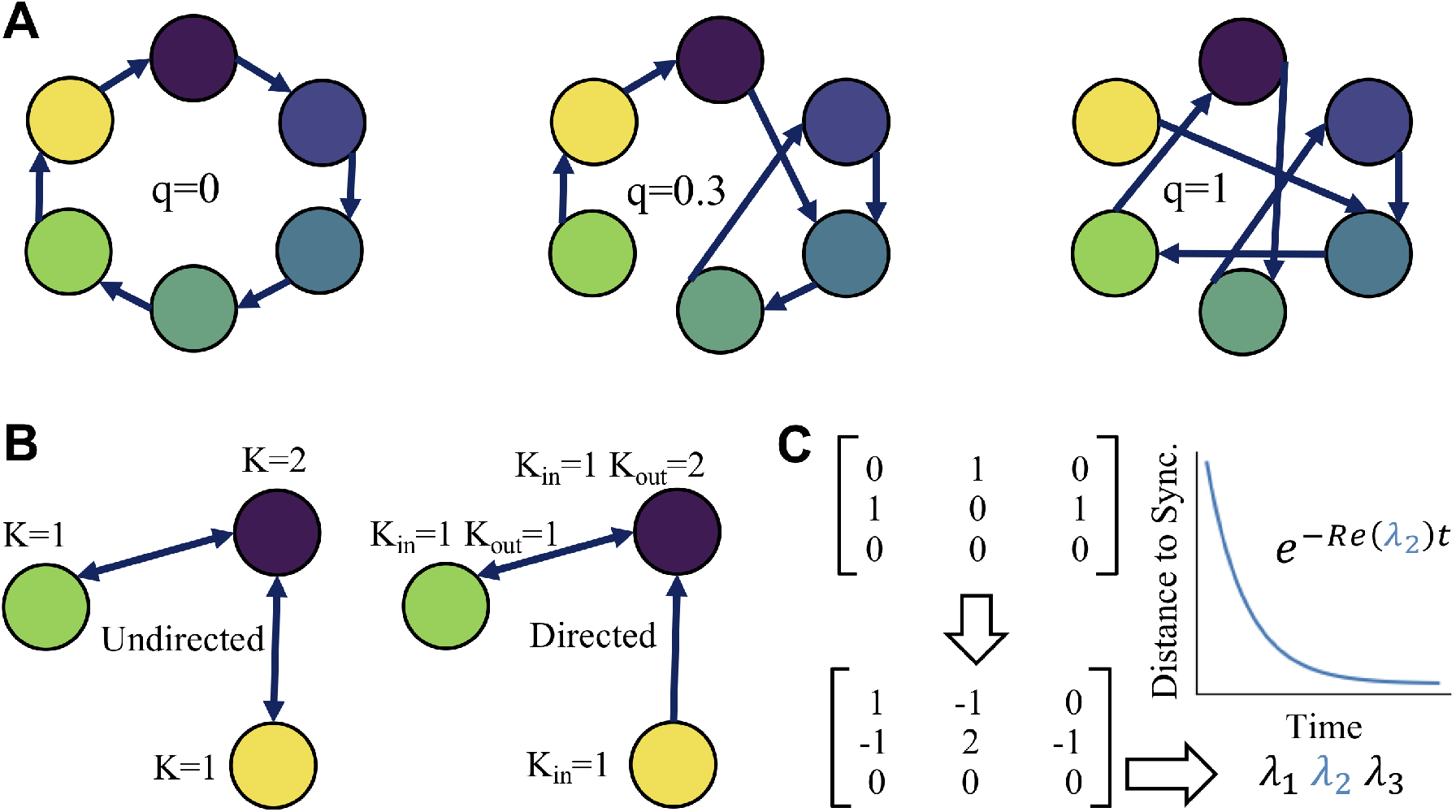
Characterizing Network Structure. Network randomness, directionality and degree are structural features that affect graph spectrum, via the Laplacian matrix, and synchronizaton speed **A.** Three networks with increasing levels of randomness: lattice network, *q* = 0; small world network, *q* = 0.3; random network *q* =1. **B.** Degree and directionality: undirected network, with *K* = *K*_in_ = *K*_out_ denoting degree for each node; directed network, with *K*_in_ denoting in-degree and *K*_out_ out-degree, for each node. **C.** Matrices associated with the directed graph in B. Counter-clockwise from top left: adjacency matrix; Laplacian matrix; eigenvalue spectrum of the Laplacian matrix; the second smallest eigenvalue dictates synchronization speed in linear dynamical systems.

Directionality (Figure 1.B) is a central feature of neural communication. Synaptic connections between two cells can be directional, as for the classic chemical synapse and rectifying gap junctions, or bidirectional, with non-rectifying gap junctions or a pair of reciprocal chemical synapses.

Degree, the total number of connections into and out of a neuron, can be subdivided into ‘in-degree’, the number of inputs, and ‘out-degree’, number of out-going synapses (Figure 1.B). As degree is not necessarily the same for all neurons in a network, networks can be characterized both by average degree and degree homogeneity, whether or not degree is the same for each neuron in the population. These dimensions are known to influence the capacity of a collection of coupled oscillators to synchronize [23, 25], or synchronizability, and mean degree has also been studied with respect to synchronization speed in certain contexts (e.g. [10]). However, the effect of degree homogeneity on synchronization speed remains, to our knowledge, unstudied.

Two matrices frequently used to characterize complex networks are the adjacency matrix (also known as the connection matrix) (Figure 1.C.i) and the graph Laplacian (Figure 1.C.ii). The adjacency matrix succinctly describes the pattern of connections in the network. It is typically a binary matrix with a 1 at the *i,j^th^* element denoting a synapse from neuron *j* to *i*, and a 0 denoting a lack of synapse. The Laplacian matrix is significant for its eigenspectrum, which is used to characterize the spectral properties of a graph in much the same way that the Fourier components characterize a time series [31]. There are several variants of the Laplacian; in this paper we use the Graph Laplacian definition, *L* = *D* – *A*, where *D* is the diagonal in-degree matrix and A is the adjacency matrix. The Laplacian’s spectrum is known to be important for dynamical processes on a graph [18, 9]. In particular, the algebraic connectivity, or smallest absolute value non-zero eigenvalue of the Laplacian, which we denote λ_2_, can be shown to determine the time scale of synchronization for many systems of coupled oscillators [3, 10] (Figure 1.D), likely due to its connection with the sparsest cut of the network’s graph [8, 41].

We note that making comparisons between directed and undirected circuits is not clear-cut. Specifically, matching number of synapses in the biological systems being modelled, the number of ‘physiological’ synapses, or matching adjacency matrix elements both seem like reasonable options but lead to a comparison of different networks. In the first case, the total number of graph edges are matched while the second case can allow mean in/out-degree to be matched. In our case, we chose to match adjacency matrix elements and mean in/out degree. We outline the implications of this in the discussion.

### Model

The simulated network was comprised of Hodgkin-Huxley neurons with parameters fit to PN pacemaker dynamics, as described in [29]. Cells were connected by linear voltage coupling to model gap-junctions with a coupling strength of *g_c_* = 5 × 10 ^−6^ mS, in accordance with experimental knowledge of PN synapses [21]. The equation for a single cell was given by

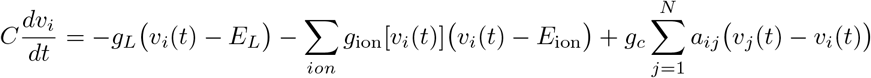

where *a_ij_* represents the *i,j^th^* element of the *N* × *N* adjacency matrix A and *N* is the number of cells in the network. For a full list of parameters see parameters and equations for canonical model (model A) in [29]

To test the effects of degree variability and directionality five matrix types were created, those with: (1) directed (rectifying) synapses and fixed (homogeneous) in/out-degree, (2) directed synapses and fixed in-degree only, (3) directed synapses and variable (heterogeneous) degree, (4) bidirectional (non-rectifying) synapses and fixed degree and, (5) bidirectional synapses and variable degree. Adjacency matrices (n=20) of each type were generated at 20 levels of connection randomness, *q* and 2 levels of mean in/out-degree, 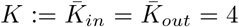 and *K* = 6 (see following), where *x* denotes mean of *x*. In/out-degree were precisely equal to mean degree in the fixed cases and were randomly distributed with the same total number of synapses and same mean degree as the fixed degree networks in the cases where in, out, or in/out, degree was variable. In the variable case in/out-degree distributions evolved from a dirac delta function, at *q* = 0, to hypergeometric distributions, when both in and out degree were variable, and binomial distributions, when only one of in or out degree was allowed to vary, at *q* = 1. Experimental work suggests the PN to be sparsely connected, with each neuron connected to roughly 5% of the total network size [21] but the degree distribution of a given cell is not known.

**Figure 2:**
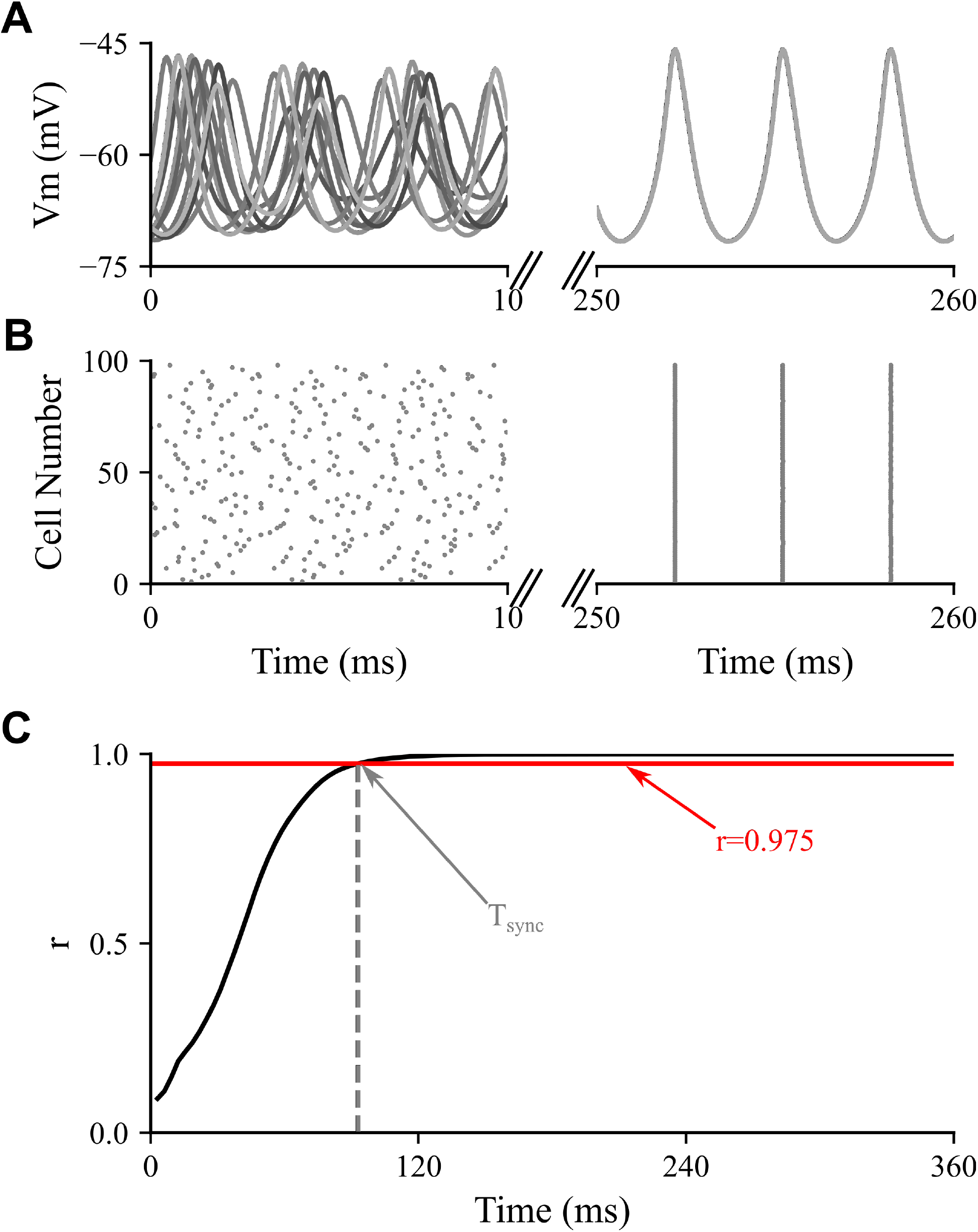
Quantifying Synchronization Speed. Synchronization times are calculated by mapping membrane voltages to phase space, calculating the Kuromoto coefficient, *r*(*t*), of the network and recording the time it takes *r*(*t*) to reach a threshold. **A.** Synchronization of membrane potentials as a function of time. **B.** Synchronization of spike times as a function of time. **C.** Example *r*(*t*) tragectory and recorded synchronization time for threshold r value of 0.975.

To assign randomness of connectivity we followed a process where each graph started as a lattice, with every node connected to K/2 of its neighbours on each side. Each connection was then randomly chosen to be removed with a probability equal to *q*. The number of synapses removed was recorded and the same number was then “rewired”, randomly, back into the network in a manner satisfying the constraints of the given graph. Beyond keeping degree fixed, where necessary, and keeping undirected adjacency matrices symmetric, we also enforced no self-loops; that is, a single neuron was not allowed to synapse with itself, and no more than one connection of a given direction was allowed between two neurons. For the fixed in/out-degree networks not every re-wiring satisfied these last two properties; to find appropriate connectivity patterns we repeatedly tested random rewirings until the two properties were satisfied. Our method is inspired by the classic method for generation of small world graphs employed by Watts and Strogatz [37, 5]. Lastly, all networks in the study were required to be complete (fully connected) as disconnected networks would never synchronize because certain neurons or sub-networks would be isolated. To ensure completeness, Tarjan’s algorithm [33] was used to locate any isolated sub-networks in each graph. The isolated components, typically single neurons, were then deleted from the network. All resulting networks contained more than 90 neurons, 98.6% contained more than 95 and 69.2% contained all 100 neurons. Notably, 100% of the directed fixed in/out-degree networks were fully connected.

Because of the randomness inherent in the network generation process, any single combination of *N, K* and *q* would generate slightly different network shapes. We thus randomly generated 20 networks for each unique combination of parameters. Initial conditions for each model pacemaker cell were chosen randomly from points on the isolated cell’s limit cycle. Simulations were run for a total of 400 ms as most networks that exhibited convergence to particular phase locked states appeared to do so over this time period. For full package specification and implementation see code availability section.

### Data Analysis and Synchronization Time Prediction

#### Quantifying Synchronization Time

We computed synchrony with a Kuramoto order parameter (*r*) [1]. Since the simulations were noiseless, spike peak could be reliably determined and used to indicate cycle time (phase = 0, 2*π*); phase increases linearly between two spike peaks. Considering the phase of each neuron as a vector on the unit circle, the Kuramoto order parameter is the magnitude of the average of these vectors, giving a value of 0 for complete asynchrony and a value of 1 for perfect synchrony.

Using this order parameter the time-to-synchronization was calculated for all simulations. A threshold value of *R* = 0.975 was used to indicate that the network was ‘synchronized’. To be sure that results were not influenced by choice of threshold, a variety of thresholds between *R* = 0.75 and *R* =1 were explored and all yielded qualitatively similar results. A bisection search in concert with interpolation was used to determine the time at which *R* = 0.975

#### Predicting Synchronization Time

To predict synchronization time from different subsets of eigenvalues from the graph Laplacian’s spectrum, we fit four different statistical models: a simple inverse function with parameter *C*(*C*/λ), inspired by previous theoretical work [3, 10], and three feed-forward artificial neural networks. The inverse function was fit by minimizing the sum of squared error (SSE) between the function and the data points (scipy.optimize.curve_fit). All artificial neural networks were implemented using PyTorch (PyTorch.org), and were composed of an input layer with as many units as the eigenvalues used for prediction in the given model, a hidden layer of 100 units and an output layer of 1 unit. Optimization used ADAM [13], with a learning rate of 0.000002, a mean square error loss function with L2 regularization (regularization coefficient = 0.05) and batch sizes of 16 samples. This training procedure was run for 60 epochs for all networks. Cross-validation on held-out data was performed, and the models did not appear to be over-fit.

### Code Availability

All simulation and analyses were performed in Python 3.7. The full set of libraries used along with code for the project is available at: https://github.com/aaronshifman/Williams_et_al_Pacemaker_Synchronization.

## Results

### Network Randomness Dictates Ability to Synchronize

The real PN robustly and rapidly returns to a synchronized, phase-locked oscillation, following behavioural modulations, without ever falling into a stable asynchronous state or exhibiting continued non-periodic or non-phase-locked behaviour [17]. However, even within the set of strongly connected graphs we have considered here, certain connectivity patterns will not exhibit this ‘synchronizability’. Thus, a necessary preliminary step was to determine which networks within this study’s parameter-space were synchronizable. For the purpose of this paper we defined synchronizable networks to be those that reached a synchronous state in less than 400 ms, as synchronization times relevant for the PN must be on the order of tens of milliseconds.

A large body of literature demonstrates the importance of network randomness for synchronization in complex networks [3, 4, 14, 12, 34], a phenomenon which we confirm in our model PN (Figure 3). Many networks with low randomness exhibited Kuromoto coefficients that did not converge towards 1 during the simulation (Figure 3.A). However, the proportion of synchronized networks (Figure 3.C) increased rapidly as a function of randomness, with all networks synchronizing during the simulation time even for relatively low *q* (Figure 3.B).

**Figure 3:**
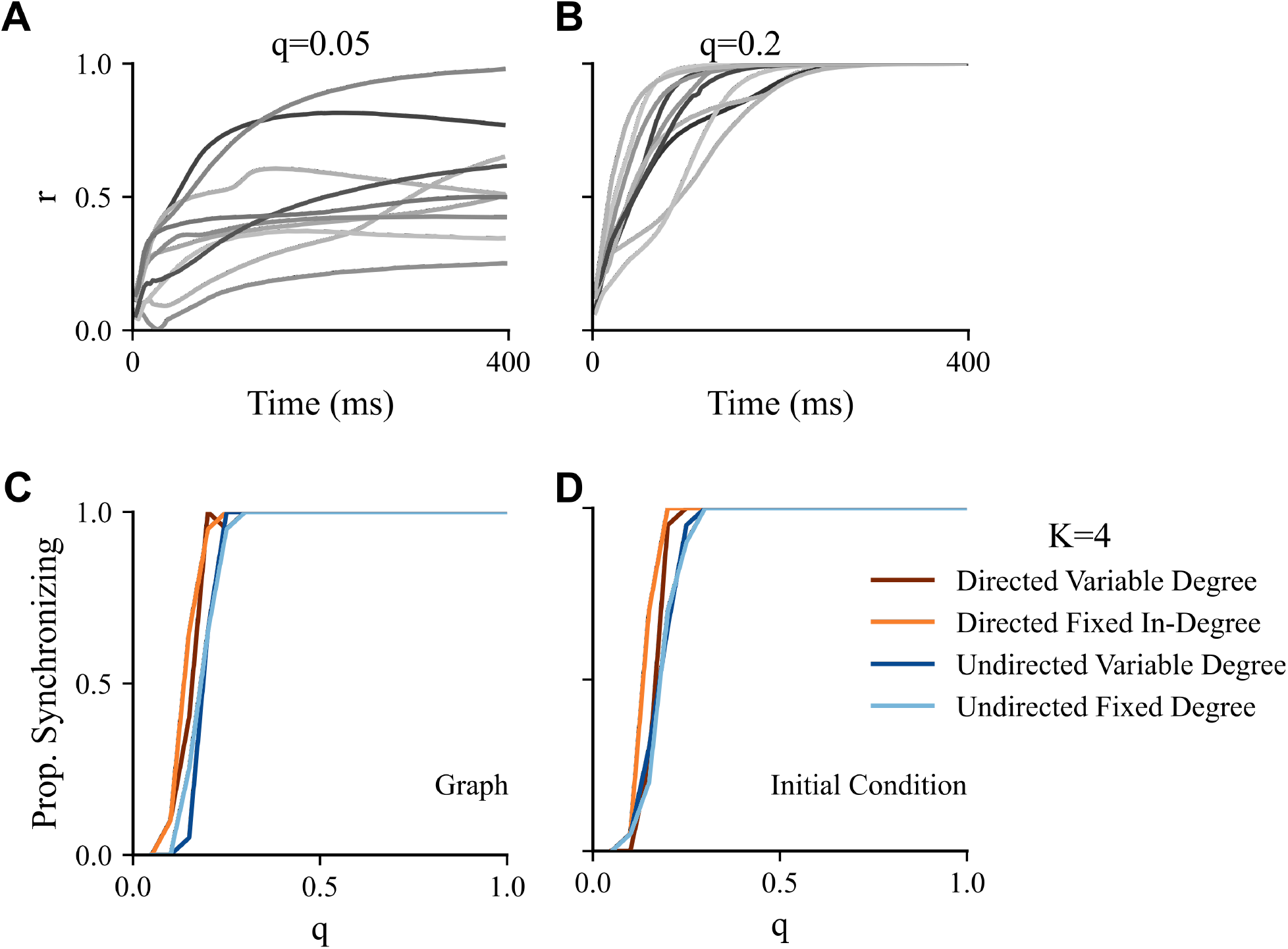
Synchronizability Depends on Network Randomness. Whether a network will synchronize depends heavily on q and weakly on the other structural parameters tested **A.** *r*(*t*) traces for *q* = 0.05; many networks do not synchronize **B.** *r*(*t*) traces for *q* = 0.2; most networks synchronize **C.** Proportion of networks in which at least 19 of the 20 tested initial conditions lead to synchrony before the end of the simulation, as a function of network randomness, *q*, for each network type (different colour denotes different network type). Note the minor differences between network types and large effect of low randomness. **D.** Proportion of initial conditions in which at least 19 of the 20 tested graph structures lead to synchrony before the end of the simulation, as a function of network randomness, *q*, for each network type (same colour scheme as C).

While randomness was the primary determinant of synchronization ability, other parameters played a small role. The network type with the highest proportion of synchronizing graphs for low randomness was directed-fixed degree, followed by directed variable degree. Both undirected network types performed equivalently across *q* values. Because this ordering closely reflects the synchronization speed related behavior we describe in the following (Figure 4), it is possible that these minor synchronizability differences could be a result of a few slow networks not converging to synchrony in the allotted timeframe, making them synchronizable in a more general sense, but not synchronizable according to our 400 ms threshold.

**Figure 4:**
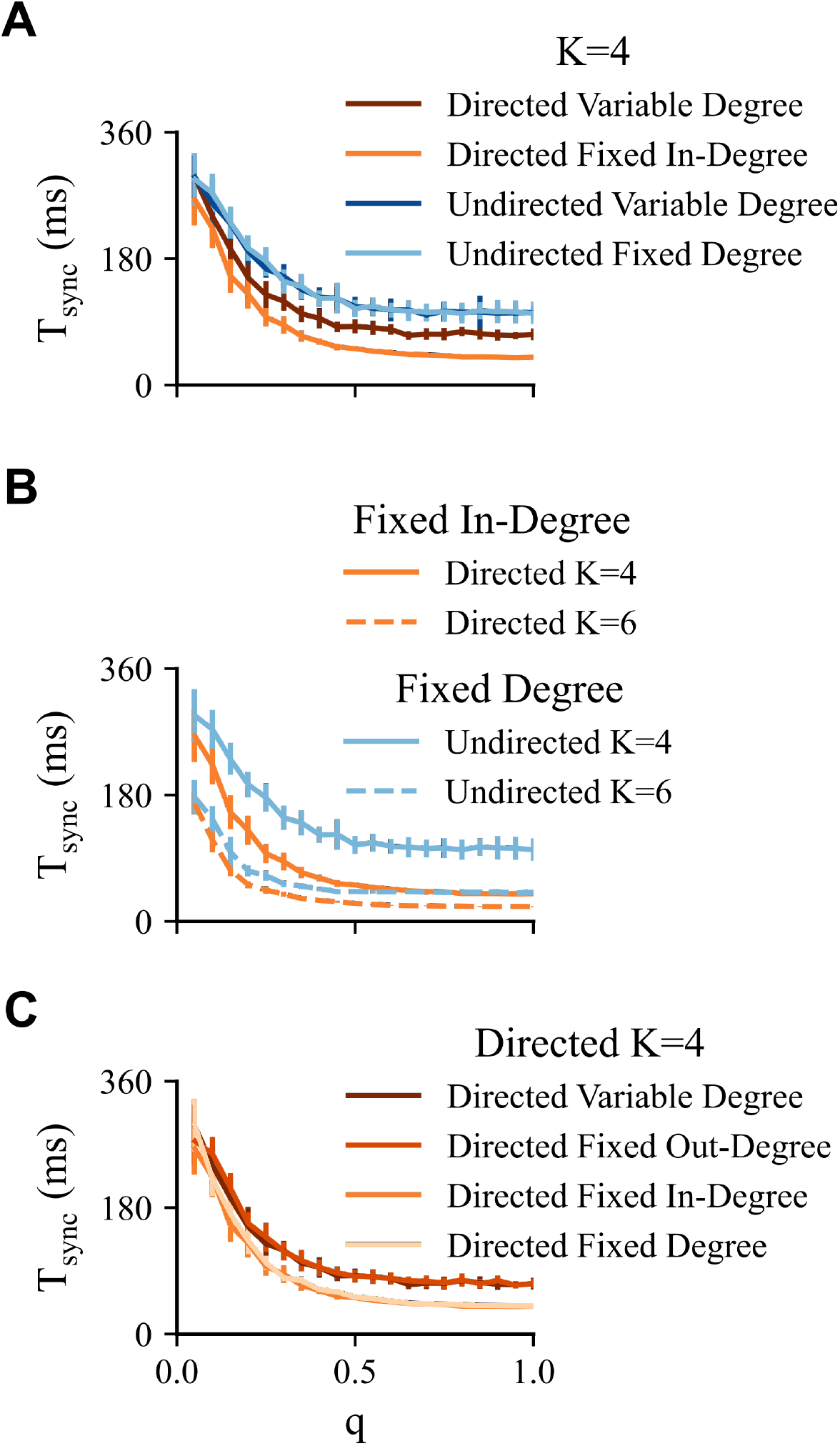
Interaction Between Degree Variability and Directedness Drive Synchronization Speed. Mean, over initial condition and graph instance, of time to synchronization, *T*_sync_, as a function of randomness, q, for different network types. Network randomness and an interaction between directedness and degree distribution exert the strongest influence on synchronization time. **A.** Comparison of four main network types; for a given mean in/out-degree, directed, fixed in-degree synchronize fastest for most *q*. **B.** Comparison of different mean in/out-degree, *K*; increasing degree speeds synchronization and reduces effect of directedness but not randomness. **C.** Comparison of fixed vs. variable degree; networks with fixed indegree synchronize the fastest. Error bars for all figures are the standard deviation of the means over initial conditions, to give a sense of the variability due to changes in graph structure.

In conclusion, the observation of greater randomness of connectivity supporting better synchronizability in our model PN is consistent with past theoretical work and suggests that the apparently high randomness in the real PN enables the robust synchronization observed. How randomness might influence the rate of synchronization is one of the questions we will address in the following section.

### Network Parameters Interact to Determine Synchronization Speed

In this section, we address the question of how connectivity parameters influence synchronization speed. Every parameter dimension explored (i.e. directedness, degree homogeneity, degree magnitude and connectivity randomness) affected time-to-synchronization, *T*_sync_, in our model PN. For the most part, the effects of these parameters were interdependent, meaning that changing multiple parameters influenced synchronization speed in a way that was different than the sum of the influence of each dimension separately. These phenomena can be distilled into four main effects on synchronization time: (i) the consistent reduction from network randomness; (ii) the benefit of directed connections (iii) an interaction between degree homogeneity and directedness; (iv) the decrease with increasing mean degree. We will address these four points sequentially.

Network randomness consistently and monotonically decreases synchronization time in our model PN (Figure 4). As in the case of synchronizability, the effect of small changes in network randomness is most notable for low q and decreases asymptotically as q increases. Past work has found this monotonic relationship between network randomness and synchronization time in a variety of coupled oscillators [10, 26]; our study extends these results to our biologically detailed PN model.

For a given value of mean in/out-degree, directed networks consistently outperformed undirected networks in our model (Figure 4.A-B). As directed synapses model rectifying gap junctions, this result suggests that rectifying electrical synapses in the PN [20] could play a functional role in the observed rapid resynchronization dynamics.

Interestingly, we observed an interaction between degree heterogeneity and directedness in determining synchronization speed. In undirected networks in-degree distribution has no effect on synchronization speed. In directed networks, changing from variable to fixed in-degree doubles the decrease in *T*_sync_ relative to undirected networks (Figure 4.A). In directed networks, degree heterogeneity can be split into heterogeneity associated with in-degree and that associated with out-degree. Notably, homogeneity only effects synchronization speed via in-degree (Figure 4.C), which is due to the fact that, in rectified networks, only the in-degree directly influences cell potentials. Degree heterogeneity increasing synchronization time is particularly intriguing given that network randomness strictly decreases synchronization time. Nishikawa et al. [25] observed opposing effects of these two forms of randomness previously, but in the case of synchronizability rather than synchronization time. They suggest this paradox could be explained by heterogeneity resulting in a smaller set of graph nodes, in this case PN neurons, monopolizing more connections and thus introducing a bottleneck in the flow of information across the network.

Finally, we find that increasing average degree not only decreases synchronization time, as expected, but also reduced the influence of other network parameters on *T*_sync_ (Figure 4.B). Thus, the key role of degree homogeneity and rectified connections in rapid synchronization may be particularly important for sparse neural networks like the PN.

In summary, if we assume that the PN must use the full toolbox of connectivity tricks to enable its rapid synchronization times, our results along with the observed sparseness of the PN suggest that the PN should have a highly random structure, with relatively homogeneous degree distribution if its synapses are indeed rectifying.

### Network Structural Parameters are Reflected in Laplacian Spectrum

Given the interdependence of individual network parameters in determining synchronization in our model PN, a natural follow up question is whether there exists a more global descriptor of network structure through which all parameters could be related to synchronization speed. The natural candidate statistic for this problem is the spectrum of the graph Laplacian, which characterizes, in a vector of eigenvalues, the structure of a complex network [18] and relates it to dynamics on the network [2, 10]. Recent work has described the smallest and largest eigenvalues of the Laplacian as a function of network parameters [9]. However, how the set of biologically relevant parameters explored in this study (e.g. synaptic rectification, degree homogeneity) relates to the full Laplacian spectrum has not, to our knowledge, been characterized; understanding how these individual parameters influence the Laplacian might enable concise quantification of PN structure in future experimental work, in a language that is directly related to synchronization dynamics.

To investigate how our parameter dimensions influence Laplacian spectrum, we plotted the mean, magnitude-ordered spectrum for different network types. Interestingly, we found that certain parameters are better encoded in distributional changes of different subsets of eigenvalues (Figure 5). Most notably, changes in average degree were better reflected in mid to large-valued eigenvalues, as can be seen from the greater separation in mean eigenvalue in the upper-right portion of Figure 5.D, while changes in our other three parameters were better encoded in the smaller and the larger magnitude eigenvalues, as is observed by the separated mean traces at left/right ends of Figure 5.A-C versus intersections in the middle portions of the plots.

**Figure 5:**
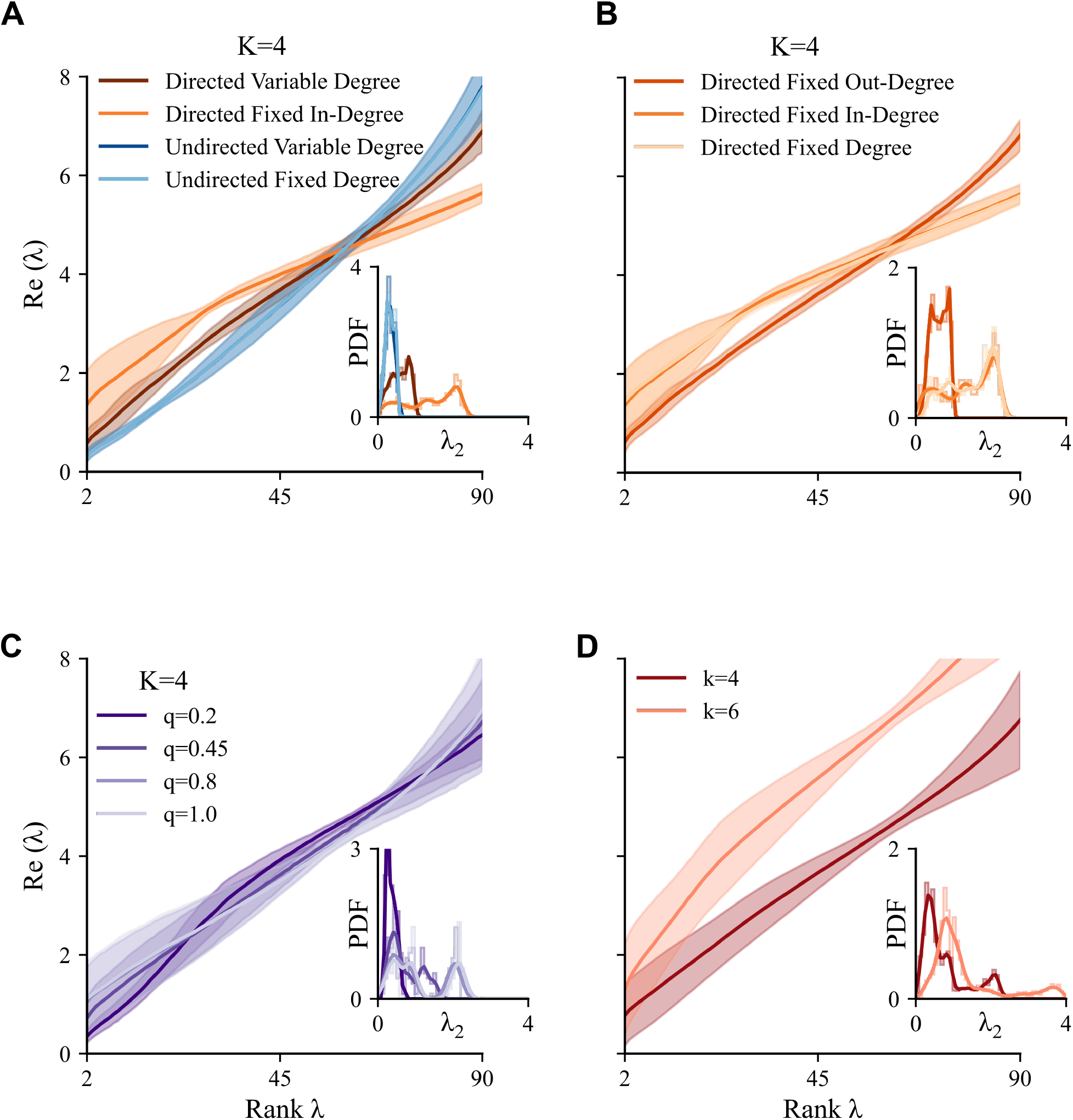
Laplacian Spectrum Encodes Network Parameters. Real part of each eigenvalue of the graph Laplacian (mean ± s.d.), for various network classes. X-axis is the rank of the given eigenvalue. Insets are distributions of the algebraic connectivity, λ_2_, cut-off at λ_2_ = 2.5. **A.** Four main network types. **B.** Directed networks with different forms of degree homogeneity. **C.** Four levels of network randomness. **D.** Two levels of average in/out-degree.

Given the significance of the algebraic connectivity, we plotted its distribution (Figure 5, insets) and found that its mean encoded network parameters in a way that is consistent with the observed relationships between network parameters and *T*_sync_. We explore this further in the following section in which the separation of parameter effects on the Laplacian spectrum motivated our analysis of *K* = 4 and *K* = 6 mean degree networks separately.

### Graph Spectrum Predicts Synchronization Speed in Model Pacemaker Network

To explore how network structure alone dictates synchronization time, we characterized the relationship between *T*_sync_ and elements of the Laplacian’s spectrum. Studies exploring dynamics in other networks of coupled neurons, and oscillating systems more generally, have observed an inverse relationship between the eigenvalue of the Laplacian with second smallest absolute magnitude and synchronization time [3, 10, 26]. However, the theory developed in these past works relies on a linearization of the system about the synchronous state, an assumption that is not valid in general.

To address this question, we began by fitting four statistical models to predict synchronization time from the eigenvalues. First, we checked how well the linearized theory [3, 10, 26] applied by using nonlinear least-squares to fit the inverse function 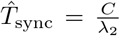, where *C* is the parameter to be fit, λ_2_ is the second smallest magnitude eigenvalue, and 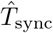 is the estimated synchronization time. Second, we tested whether some other relationship might better predict synchronization time from λ_2_ by fitting a feed-forward artificial neural network to predict synchronization time from this eigenvalue. Finally, we fit two other artificial neural networks to determine *T*_sync_ from the first 20 non-zero eigenvalues and the first non-zero 90 eigenvalues of the Laplacian, respectively. This was done to determine whether more information about synchronization dynamics might be contained in a larger subset of eigenvalues than just λ_2_ alone. Motivated by the qualitative similarity between mean eigenvalues within networks of a given degree (Figure 5.A-C) versus the distinct differences in mean spectrum between average degree (Figure 5.D), we fit the four models separately to the *K* = 4 and K = 6 networks.

We find that very little extra information is contained in the spectrum beyond the algebraic connectivity, regardless of mean degree (Figure 6.B). However, the effectiveness of the linearization analysis-derived relationship, the inverse function, at relating algebraic connectivity to synchronization speed depends on average degree. For *K* = 4, arbitrary function approximation via the artificial neural network is able to better predict the nonlinear relationship between eigenvalue and synchronization time, while for *K* = 6 the inverse function predicted by linearization performs at essentially the same level of error as the artificial neural net. We hypothesize that the inverse function doesn’t fit synchronization time on the lower value of *K* as well on account of a longer period of nonlinear dynamics, far from the synchronous fixed point, as observed previously in other systems of coupled oscillators [43, 34]. It bears mentioning that the inability of the artificial neural network to detect extra information in the Laplacian outside of that contained in λ_2_ could be the result of the non-convex optimization process being stuck in a local minimum or the true function being outside the set of functions that the tested artificial neural network could model, though this seems unlikely.

**Figure 6:**
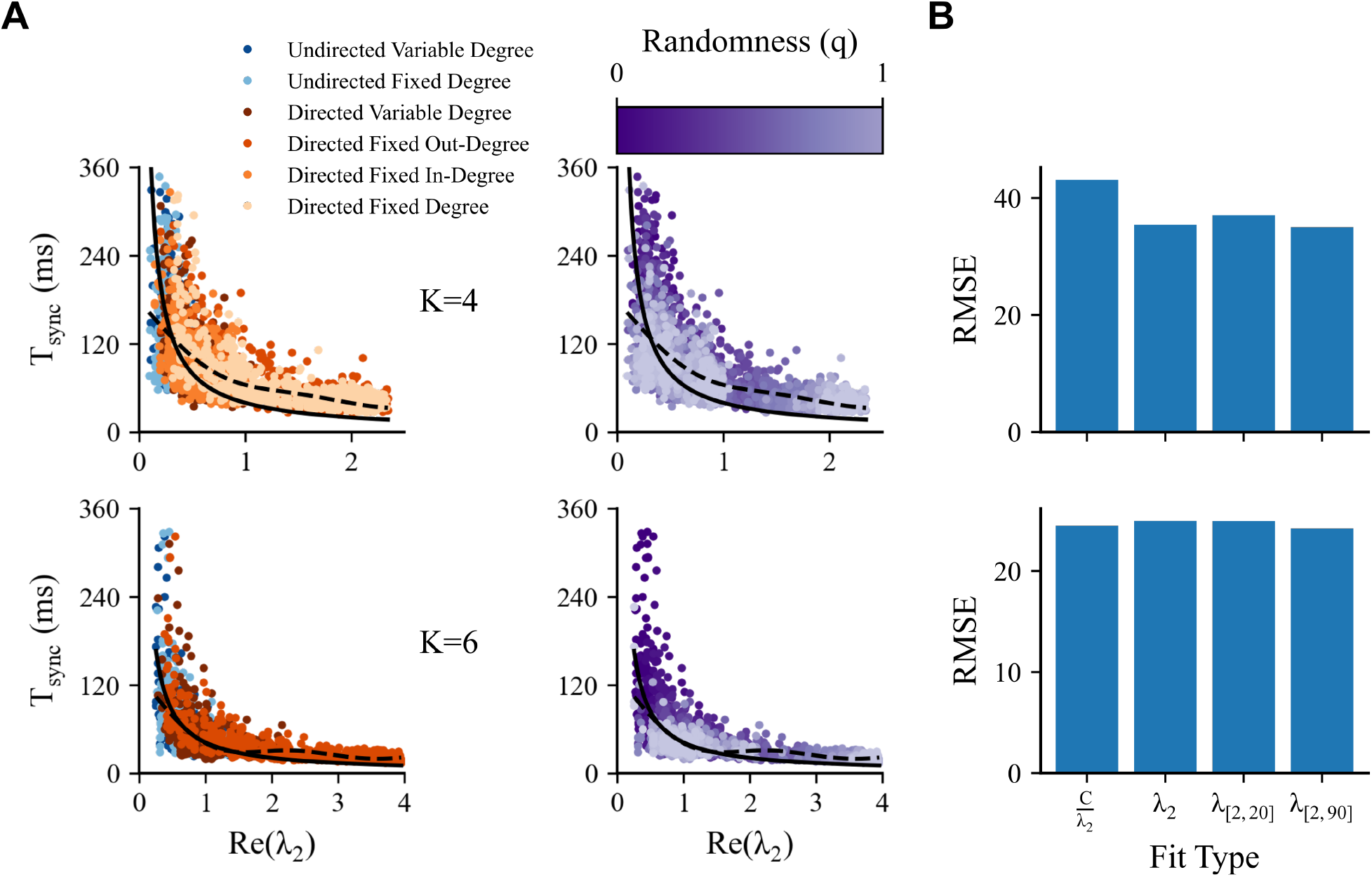
Algebraic Connectivity Predicts Synchronization Time in Model Pacemaker. **A.** Scatter plots of synchronization time versus algebraic connectivity, λ_2_, coloured according to network type (left column) and network randomness (center column). Top row gives data for mean in/out-degree *K* = 4 networks (note high variability in scatter plots) and bottom gives data for *K* = 6 (note comparatively lower variability). Plotted lines are prediction of models *C*/λ_2_ (solid) and an artificial neural net fit on λ_2_ (dashed). **B.** Root Mean Square Error (RMSE) for each model on validation (held-out) dataset. Left to right: *C*/λ_2_, *FNN*(λ_2_), *FNN*({λ_2_, λ_3_,…, λ_20_}), *FNN*({λ_2_, λ_3_,…, λ_90_}).

For a given mean degree, the different network parameters are all related to synchronization time through the algebraic connectivity, with ‘‘slower” parameterizations, like undirected networks, falling to the left in the scatter plot (Figure 6.A.i and B.i) and ‘faster” parameterizations, like directed fixed in-degree networks, falling to the right. The banding of the scatter plots in Figure 6.A is due to the full range of q values being exhibited in undirected networks and again in directed networks, and provides an alternate view of the result from Figure 4: that, for a given mean degree, directed networks are capable of matching undirected networks in synchronization time for much lower network randomness.

## Discussion

In this study, we investigated the role of network randomness, synapse directionality and degree homogeneity on synchronization speed in a model PN. We found that, when mean in/out degree rather than number of phenomenological synapses is matched, random, directed networks with homogeneous degree synchronize the fastest. Moreover, we found network structure, as characterized by the second smallest eigenvalue of the Laplacian, to be predictive of synchronization speed, a relationship suggested by previous studies [10, 3]. However, the accuracy with which the inverse relationship between eigenvalue and synchronization time, predicted by these studies, depended on the mean degree of the PN model. These results have implications both for the PN itself and for neural circuits more generally.

For the PN, our work suggests that the characteristic rapid synchronization times could be driven by high randomness of connectivity and, in the case of rectifying PN gap junctions, a less-variable degree distribution. These findings provide a quantitative context for past experimental work which has suggested random connectivity and rectifying synapses to be features of the PN. We also suggest that the homogeneity of degree should be considered in future experimental studies as a new connectivity-related marker for rapid synchronization dynamics. Interestingly, our modelling demonstrates that these properties lead to synchronization times that are close to, but still fall short of, the 5-period synchronization times sometimes observed in the PN [17]. There are several potential mechanisms that could close the gap between observed PN synchronization time and times seen in this study. Our work has focused on synchronization of neurons from initial conditions (random phases) on the neurons’ limit cycle; how the resulting dynamics differ when biological desynchronization (e.g. synaptic input) drives neurons off their limit cycles is not clear. Although it is expected that pacemaker cells alone dictate synchrony in the PN, experimental work has not yet fully discounted the role of other cells, e.g. relay cells [20, 6], in synchronization dynamics. Finally, it has also been suggested that electrical field interactions, i.e. ephaptic coupling, could play a role in boosting synchronization speed in the PN [29, 17]. Determining the precise combination of additional phenomena that enable the PN to synchronize so rapidly presents an exciting direction for future experimental research.

Another important direction for future research is to further investigate synchronization speeds in directed versus undirected networks. In this study we compared networks where the number of non-zero elements in the graph adjacency matrix were matched. This seemed logical because it allowed us to match in/out-degree between directed and undirected graphs. However, because a single undirected synapse is represented in the adjacency matrix as two directed synapses of opposing direction, this paradigm meant that the directed networks had twice as many phenomenological synapses as the undirected kind. Future work is necessary to determine if directed networks still synchronize faster than undirected when one matches phenomenological synapses; e.g. if one compared undirected networks with mean in/out-degree of 8 with directed networks of mean in/out-degree of 4. While our comparison (see Figure 4.B) of *K* = 4 directed with *K* = 6 undirected networks suggest that undirected networks of *K* = 8 could synchronize more rapidly, a fair comparison necessitates matching total conductance between graph types and this remains to be done.

An limitation of this study is with regards to network generation: we have been content to use graphs that satisfy our parameter combinations while, empirically, appearing to exhibit an otherwise random sampling from the space of potential graph adjacency matrices. A more mathematical analysis of the graph generating algorithms is warranted to provide context for the results of this study. By exploring the statistics of these methods, such as the probability distributions on adjacency matrix elements, one could answer questions such as why the generation of directed, fixed in/out-degree networks almost always resulted in fully connected graphs, and whether the study results are quite general or may be influenced by an under/over-sampling of the probability space associated with a given graph type.

Finally, our results have implications for the study of synchronization phenomena outside the PN; namely, the illumination of network structure-related markers of rapid synchronization dynamics. The ability to locate neuronal circuits underlying rapid spike-time synchronization may improve detection of brain regions susceptible to epileptic seizures [38, 24]. Our work provides both qualitative descriptors, e.g. directed connections with homogeneous degree, and a quantitative approach, artificial neural network fitting to the algebraic connectivity, for predicting how quick a given network will synchronize via its eigenvalues. In particular, this study suggests that there may be more predictive information to be gained from the algebraic connectivity, in sparse networks, than previous studies employing linearized methods might have implied.

## Acknowledgments

This work was supported by an Undergraduate Student Research Award and Canada Graduate Scholarship - Masters (CGS-M), from the National Sciences and Engineering Research Council of Canada (NSERC), and an Ontario Graduate Scholarship (OGS) from the Ontario Student Assistance Program to EW, an NSERC Alexander Graham Bell Canada Graduate Scholarship (CGS-D) to AS and an NSERC Discovery Grant to JL (05872).

